# Temporal gradient characteristics of automatic neural representation for orthographic consistency in Chinese characters

**DOI:** 10.1101/2022.07.25.500986

**Authors:** Jianyi Liu, Tengwen Fan, Yan Chen, Jingjing Zhao

## Abstract

Visual word processing involves the automatic decoding of orthographic, phonological and semantic information. The mappings of these information in a writing system comprise an important feature of visual words: orthographic regularity and consistency. Recent electroencephalography (EEG) studies have investigated the automatic processing mechanism of orthographic regularity (i.e., words vs. pseudowords) among visual words. However, the automated mechanism and temporal dynamics of different types of orthographic consistency effects (e.g., orthographic positional consistency, orthography-to-phonology consistency vs. orthography-to-semantics consistency) have never been studied. This study explored automatic neural representation for orthographic consistency effects in visual words and its temporal dynamics through an oddball paradigm. Three types of oddball sequences were designed with Chinese characters as stimuli, including consistent Chinese characters as standard stimuli and three types of inconsistent characters (orthographic positional inconsistent, orthography-to-phonology inconsistent vs. orthography-to-semantics inconsistent) as deviant stimuli, respectively. Significant visual mismatch negativity (vMMN) activities were observed in all three types of inconsistent characters, which suggests automatic processing caused by orthographic consistency violations. Time-resolved representational similarity analysis (RSA) further revealed that there are different temporal dynamics of automatic neural representations for the three types of consistency features. The representation of positional consistency emerged earlier within an independent time window, while the representation of phonetic and semantic consistency emerged later, and partially overlapped. These findings provide novel insights for the temporal gradient characteristics of automated representation structure of orthography consistency information.

## 1 Introduction

Humans inherit the wisdom of their predecessors through reading. Rapid and automatic decoding of visual words is a prerequisite to acquire such a core cultural skill. The content of visual word decoding includes both the word form (i.e., word length, letter order, and radical position; Liu et al., 2021), as well as high-level lexical information (i.e., phonology and semantics; Carreiras et al., 2014). In addition to its own statistical properties (e.g., frequency and/or legality of consonant doublets in alphabetic language; the frequency of a radical occurs in a given location within characters in Chinese), visual words from information have a stable statistical mapping with nonvisual lexical information (i.e., orthography-to-phonology (O-P) mapping and orthography-to-semantics (O-S) mapping; Zhao et al., 2018). This probability information, collectively known as orthographic regularities, has an important function in the neural specialization for print (Zhao et al., 2019; Tong et al., 2020).

Consistency of structural or positional, phonetic, and semantics are considered to be key aspects for orthographic regularities (Zhao et al., 2014; He et al., 2017; Tong et al., 2020). This means that the probability that the properties of a particular feature (i.e., pronunciation) of a sub-lexical element in the current word are the same as those of corresponding properties, among other words composed by identical elements. The classic example of consistency effect on word processing is that words with an orthographic body reliably paired with a particular phonological rime (e.g., “consistent” words, which refers to a group of words with a high probability in a particular word set, such as pill, mill, and still) are read aloud faster and more accurately than words that have inconsistent body-time correspondences (e.g., pint; Glushko, 1979; Jared, 2002).

The consistency features of the Chinese writing system are mainly placed among two elements called radicals, one that provides information about how that character is pronounced (phonetic radical), and the other providing information about its meaning (semantic radical). These phonetic-semantic compounds are also commonly referred to as “phonograms” (Zhao, 2014; Myers, 2019). For example, in Chinese phonograms 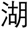hu:2/ (to lake), the phonetic radical 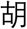hu:2/ reveals its pronunciation (/hu:2/), while the semantic radical 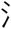 indicates its semantic category (water-related concept). In reality, however, many radicals clearly deviate from positional and lexical consistencies to varying degrees. The obvious feature of the positional distribution of phonograms is that the majority of them are left-right horizontally structured characters (around 69%; according to Myers, 2019). This, in turn, implies the presence of other structures (positional inconsistent characters). At the lexical level, approximately 35% of phonograms are phonetic inconsistent characters that differ from the common pronunciations of other characters made up of the same phonetic radicals. Furthermore, 12% are semantic inconsistent characters that differ from the common meaning categories of additional characters that are made up of the same semantic radicals (Shu et al., 2003).

The consistency effect in Chinese phonograms has been demonstrated through a series of reading aloud experiments (Fang et al., 1986). Consistent characters were named significantly faster than inconsistent characters (Lian, 1985; Lee et al., 2005). Furthermore, studies that use the artificial orthographic learning paradigm validated the significant impact of consistency on character recognition (i.e., better performance with high positional consistency) and generalization for Chinese orthographic learning (He et al., 2017; Tong et al., 2020). With regards to neural mechanisms, previous fMRI studies have reported greater activation in the left inferior frontal gyrus, the left temporoparietal (inferior parietal gyrus and supramarginal gyrus) region, and the left temporal–occipital junction when naming inconsistent phonograms compared to consistent ones. Additionally, several event-related potential (ERP) studies have validated the consistency effects of phonograms within the time windows of N170, P200, and N400 components in the homophone judgment task, lexical decision task, and delayed naming task (phonological consistency: Lee et al., 2007; Hsu et al., 2009; Yum et al., 2014; semantic consistency: Hsu et al., 2021). Similarly, based on the artificial orthographic learning paradigm, several studies have validated the modulation effect of consistency based on the corresponding ERP components evoked by new learned characters (semantic consistency: Tong et al., 2020b; positional consistency: Tong et al., 2020c).

All of these studies on the consistency of phonograms utilized explicit cognitive tasks, so that they are limited to explaining the intentional processing mechanisms of Chinese characters. However, the effortless reading comprehension that is usually experienced by skilled readers relies more on the rapid and automatic retrieval of orthographical, phonological, and semantic information. The automatic or implicit processing of written words can be reflected using visual mismatch negativity (vMMN) (for reviews see Czigler, 2014). The vMMN is usually evoked through the use of a passive (employ attention-demanding primary tasks to limit the attention resources of participants) oddball paradigm in which a standard visual stimuli type is commonly presented to the observer, while a deviant type is infrequently presented. Previous vMMN studies initially focused on automatic discrimination of physical features, including color (Czigler et al., 2002; Thierry et al., 2009), line orientation (Astikainen et al., 2008; Czigler and Pato, 2009), and spatial frequency (Maekawa et al., 2005; Sulykos and Czigler, 2011). Subsequent studies have discovered that vMMN can also reflect the automatic mechanism of more abstract properties, which include orthographical, phonological, and semantic processing of visual words (Shtyrov et al., 2013; Wang et al., 2013; Wei et al., 2018; Hu et al., 2020). However, none of these prior studies examined whether automatic consistency processing also occurs when a human observes visual words.

An important limitation of these previous studies that investigated the neural mechanisms behind orthographic regularity, including consistency, is that they are not able to simultaneously examine the interactive representations of orthographic, phonological, and semantic regularities (Tong et al., 2020b; Zhao et al., 2019; Liu et al., 2021). With the alphabetic languages, this limitation is caused by the nature of these languages, which are substantially more asymmetrical between the two mappings, with highly consistent O-P mapping and arbitrary O-S mapping (Zhao, 2014). However, the Chinese phonograms that convey a relatively symmetric statistical structure between the O-P mapping and O-S mapping provide an opportunity to explore these relevant issues. In more detail, phonograms that are inconsistent in any one dimension may be consistent in the other dimensions (i.e., 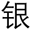 /yin/, to silver, is inconsistent for common pronunciation: /hen/, but inconsistent with the meaning category of 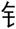 as metal-related concept, and has a common radical position). Therefore, we can manipulate the consistency of these three dimensions simultaneously in a Chinese character.

The present study seeks to understand the automatic processing mechanism of consistent features in the visual-orthography, phonetic and semantic dimensions. On this basis, we also tried to investigate the temporal dynamic relationship between neural representations of consistency across different dimensions. In order to accomplish this, we carefully selected several consistent and inconsistent phonograms (see Material section) and set the inconsistent phonograms of certain dimensions as the deviant stimuli that is present in each of the three oddball blocks with consistent phonograms as the standard stimuli. Furthermore, in order to avoid repetition effects due to refractoriness, we also set an equal probability block in which all types of inconsistent and consistent phonograms were presented with the same probability. The participants’ brain responses to the deviant were compared to responses to the control (the corresponding deviant in equal probability blocks) in order to determine whether the consistency deviations were able to evoke vMMN. Time-resolved representational similarity analysis (RSA) was carried out to track representations for inconsistent information (consistency violations) for all dimensions (Kriegeskorte and Kievit, 2013; Cichy et al., 2014).

## 2 Materials and methods

### Ethics statement

All participants gave oral and written, informed consent in accordance with procedures that were approved by the ethics committee at the School of Psychology, Shaanxi Normal University (Approval No. HR 2021-05-002). The protocols adhered to the Declaration of Helsinki.

### Participants

A total of 45 right-handed (via self-report) healthy participants (mean age = 18.04, SD = 0.80; 35 females), with normal or corrected-to-normal vision took part in the study. Participants were recruited from the undergraduate and postgraduate student population at Shaanxi Normal University and were paid 60 RMB for their participation. All participants reported no speech or hearing problems and had no prior history of neurological or psychiatric abnormalities.

### Material

There were four different sets of Chinese phonograms selected. Each set contained four characters (each row in Figure 1A). In Chinese, a character is generally made up of a semantic radical and a phonetic radical, known as “phonograms”. For example, the character 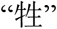 consists of a semantic radical 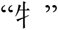 and a phonetic radical 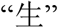. Each phonogram set in the present study has a fixed phonetic radical and four different semantic radicals (Figure 1A). In the second row in Figure 1A, for example, the phonetic radical in all characters is 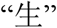, while the semantic radicals are 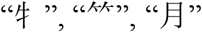, and 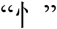, respectively.

**Figure 1.**
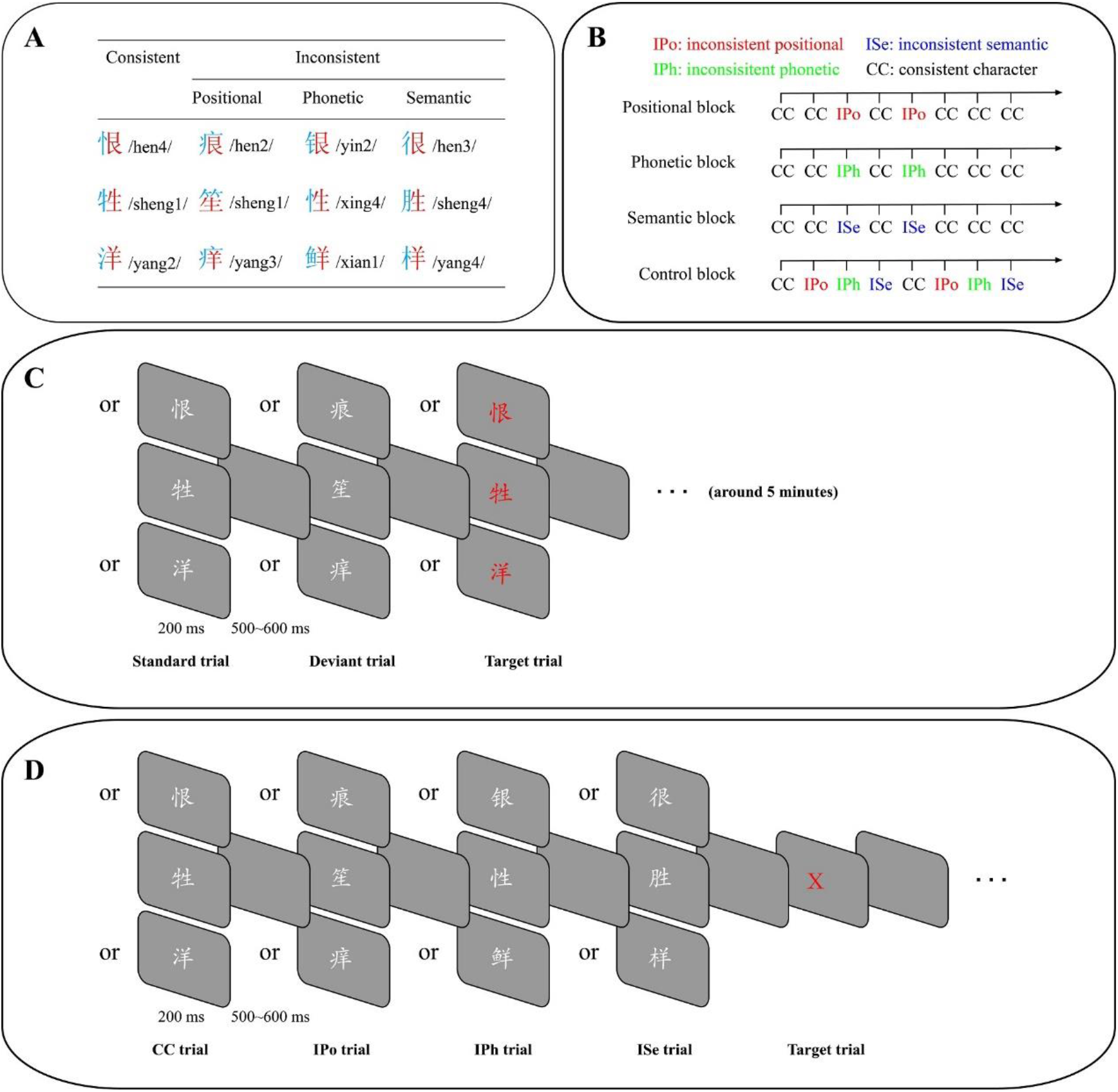
Illustration of the experimental procedure. (A) Details of the selected Chinese phonograms. (B) Examples of presentation settings for consistent and inconsistent phonograms in different blocks. (C) Schematic depiction of the color-change judgment task in the oddball block. (D) Schematic depiction of the color-change judgment task in the control block. Abbreviations: CC consistent character, IPo inconsistent positional character, IPh inconsistent phonetic character, ISe inconsistent semantic character.

Semantic radicals are generally on the left side of characters, and phonetic radicals are on the right, which is the main visual-orthography rule in the majority (63%) of phonograms (Myers, 2019). Semantic radicals usually provide semantic information of characters, while phonetic radicals provide phonological clues. For example, in the Chinese character 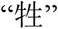, the semantic radical 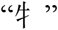 is on the left side, while the phonetic radical 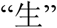 is on the right. The meaning of character 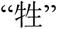 is “domestic animals”. This means that it can easily be speculated from the semantic radical 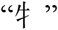, which refers to “cattle”. Phonologically, the pronunciation of 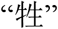 is “sheng”, which is also highly consistent with the sound of the phonetic radical 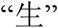 (sheng). Characters like 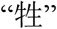 are consistent characters. However, there are some characters (i.e., inconsistent characters) that do not follow the orthographical (positional), phonological and/or semantic rules (e.g., 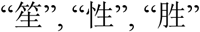).

Accordingly, these 12 Chinese characters were then divided into three consistent and nine inconsistent characters. The nine inconsistent characters were further divided into three categories, including inconsistent positional (IPo) characters, inconsistent phonology (IPh) characters and inconsistent semantics (ISe) characters. Each category has three characters that come from three different sets. In other words, all four categories (i.e., 1 consistent and 3 inconsistent categories) have the same number of characters and the same phonetic radicals (see Figure 1A). The inconsistent categories differ from the consistent category with regards to visual-orthography, phonology and semantics, respectively.

Specifically, compared to the consistent category, IPo characters differ with regards to the structure of characters. That is, phonetic radicals appear on the right side of a consistent category. On the contrary, phonetic radicals in the IPo characters are in the less common position in a character. For example, phonetic radial 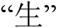 posits at the right side of a 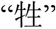, but it is at the bottom of the IPo character 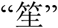. Among all characters that are 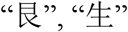 and 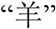, they act as phonetic radicals. The probabilities that 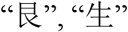 and 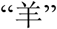 appear on the right side of characters are 72.22%, 50.00% and 72.73%, respectively. On the other hand, the percentages of the character positions for 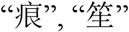 and 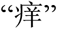 are 5.56%, 25%, and 18.18%, respectively (Supplementary Table 1). Furthermore, the position of phonetic radicals in IPh and ISe categories are on the right side, the same as consistent characters.

Similarly, IPh characters differ from consistent characters (CC) with regards to phonological consistency. The pronunciations of CC are high in phonological consistency of corresponding phonetic radicals, while the phonological consistency is low in IPh characters. The phonological consistency is defined as the proportion of a specific pronunciation among all characters that adopt the same phonetic radicals. High consistency refers to the pronunciation of a regular character that is the main pronunciation of all Chinese characters utilizing the specific phonetic radical. In contrast, the pronunciation of IPh characters is a rare sound that corresponds to phonetic radicals. For instance, the pronunciation of 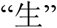 and the regular character 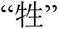 are /sheng/, which is the same as most characters that contain 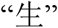 (e.g., 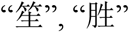). However, pronunciation of the IPh character 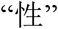 is /xing/, not “/sheng/”. The phonological consistency is 0.3874-0.9207 for all three CC and is 0-0.2831 for the three IPh characters (see Supplementary Table 1 for details). In addition, ISe and IPo characters have the same pronunciation as regular characters (Supplementary Table 1).

Finally, the ISe characters differ from the consistent ones with regards to the transparency of the semantic radical. The CC is high in the transparency of semantic radicals, while the transparency in ISe characters is low. The transparency is defined as the connection between the meaning of the semantic radical, and the meaning of the corresponding character (Shu et al., 2003). That is, semantic radicals of CC can reflect the meaning of corresponding characters. However, the meanings of the ISe characters cannot be speculated from the semantic radicals. For example, the semantic radical 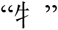 (cattle) is related to the meaning of 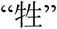 (livestock). In contrast, the meaning of ISe character 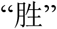 (victory) is much different from that of the corresponding semantic radical 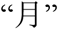 (moon). The same as CC, characters in the IPh and IPo categories are high in the transparency of the semantic radical.

### Procedure

The experimental procedure consisted of three oddball blocks and a control block. One of the three categories of inconsistent characters (each category contains three specific characters), in turn, served as the deviant stimuli (dev; probability of occurrence p = 0.25; the number of presentations is divided equally among the three specific characters) across different oddball blocks (each block contains 420 trials), while the consistent characters (containing three specific characters) served as standard stimuli (std; p = 0.75). In the control block (contains 480 trials), the probability of inconsistent (three categories; named equiprobable control stimuli) and consistent characters were the same (p = 0.25). Within each block, the trial order was fully randomized, and the order of oddball blocks was also randomized while the control block was implemented at the beginning. For each individual trial, the stimulus was presented for 200 ms, and then a gray image was inserted, lasting for 500 ~ 600 ms, at a random time between trials (Figure 1B). Moreover, the color of the characters may change from white to red at random during some trials (target; p = 0.1; may appear on the standard stimuli of the oddball blocks as well as all stimuli of the control block). The task throughout the experiment was to pay attention to whether the color was changed, as well as to press a button after each change. The deviant stimuli in the oddball blocks would not appear twice in a row, and the target only appeared after one standard trial. Participants sat comfortably in an armchair at a distance of 60 cm from the screen, and were given a break for each block that they completed. Using the E-prime software, the images of words were presented within the central visual field (visual angle: horizontally = 2.5°; vertically = 3.8°).

### Behavioral analysis

A participant’s response was counted as a hit if the button was pressed for less than 700 ms after the character color changed. Otherwise, the response was counted as a false alarm. Hit and false alarms (FAs) rates during the target detection task were analyzed in order to evaluate the degree of attention of each of the participants.

### EEG recording and preprocessing

Electroencephalography (EEG) signals were recorded through the use of a 64-channel amplifier (ANT Neuro EEGO, Inc.) that was mounted on an electrode cap according to the international 10–20 system. The online reference electrode during the data collection was CPz. The EEG data was digitized at a sampling rate of 1000 Hz, and impedances were kept below 20 kΩ during the experiment.

Offline preprocessing was conducted utilizing the EEGLAB (Delorme and Makeig, 2004) toolbox in MATLAB. EEG data were referenced to an average reference (excluding the EOG, M1 and M2 channels), that were filtered between 0.1 and 40 Hz through the use of a built-in EEGLAB function eegfilt. This utilized a two-way least squares FIR (finite impulse response filtering). The epochs were created from – 300 ms pre-stimulus to 700 ms post-stimulus for each trial, and the baseline was corrected using the first 200 ms. Epochs with any excessive movements or eye blinks (i.e., voltage exceeding ± 75 μV) were automatically rejected.

### Visual mismatch negativity (vMMN) analysis

The differential wave of characters with different inconsistent categories was obtained by subtracting the ERPs of the corresponding deviant stimuli from the ERPs of the corresponding equiprobable control stimuli. This method allows the comparison of ERPs that are evoked by the deviant of the oddball sequence to the ERPs that are evoked by physically identical stimuli from a sequence without any particular frequent (standard) stimulus (Stefanics et al., 2014).

The method of equal probability control was suggested in order to deal with repetition effects due to refractoriness that was assumed to be present in the deviant, minus standard activity that was obtained in classical oddball paradigms (Schröger and Wolff, 1996; Jacobsen and Schröger, 2001). Activity considered as “genuine” vMMN (i.e., vMMN without stimulus-specific refractoriness effects superimposed) emerges when the oddball deviant evokes a larger negativity than the control stimuli (Stefanics et al., 2014). Next, a cluster-based permutation test was utilized to search “genuine” differential activity between the ERPs of deviant and equiprobable control stimulus (Maris and Oostenveld, 2007). We conducted this analysis through the use of the Fieldtrip toolbox (Oostenveld et al., 2011) in MATLAB. We developed grand-averages of differential waveforms across two regions of interest (ROI) that correspond to the left (P7, PO7, O1) and right (P8, PO8, O2) posterior occipital-temporal electrodes (the electrodes were selected based on previous studies, e.g., Hu et al., 2020; Kovarski et al., 2021). For each time point (within 0 – 700 ms) at left or right electrodes, the clusters were formed through two or more neighboring time points whenever the *t* values (obtained by two-tailed *t*-test) exceeded the cluster threshold (0.025). The number of permutations was set to 5000, and the corrected significance level was set to 0.05. That is, when the clustering level error probability of a cluster was less than 0.05, then it was considered that there were significant effects in the corresponding period (i.e., effective vMMN activities were identified). We will report the temporal range of the significant negative clusters and their mass for each inconsistent category (the sum of *t* values in a cluster).

### Representational Similarity Analysis (RSA)

Representational similarity analysis was also conducted on the EEG data to determine the neural organization of consistency representations. First, we developed time-resolved neural representational dissimilarity matrices (RDMs), which reflected the pairwise dissimilarity of the characters’ brain representations across the processing times. Next, we compared organization of the neural RDMs to a set of predictor RDMs, which allowed us to capture different consistencies on which the characters were similar or dissimilar.

#### Neural dissimilarity

The representational dissimilarity matrix (RDM) was then calculated in order to quantify the degree of dissimilarity across neural activities evoked by different inconsistent characters within oddball blocks. The EEG RDMs were constructed separately for each participant through the use of NeuroRA toolbox (Lu & Ku, 2020). Neural RDMs were created for 70 time bins of 10 ms width each that ranged from 0 to 700 ms, relative to stimulus onset. All the following analyses were done separately for each of these time bins. For each time bin, we extracted a response pattern that was prevalent across 10 time points (covering 10 ms at 1000 Hz) and 61 electrodes by unfolding these data into a 610-element vector. In order to obtain the series of RDMs, we first averaged all available trials across for each inconsistent character, and then correlated the response vectors for each pairwise combination of characters. These correlations were subtracted from 1 and then entered into a 9-by-9 RDM (Figure 2).

**Figure 2.**
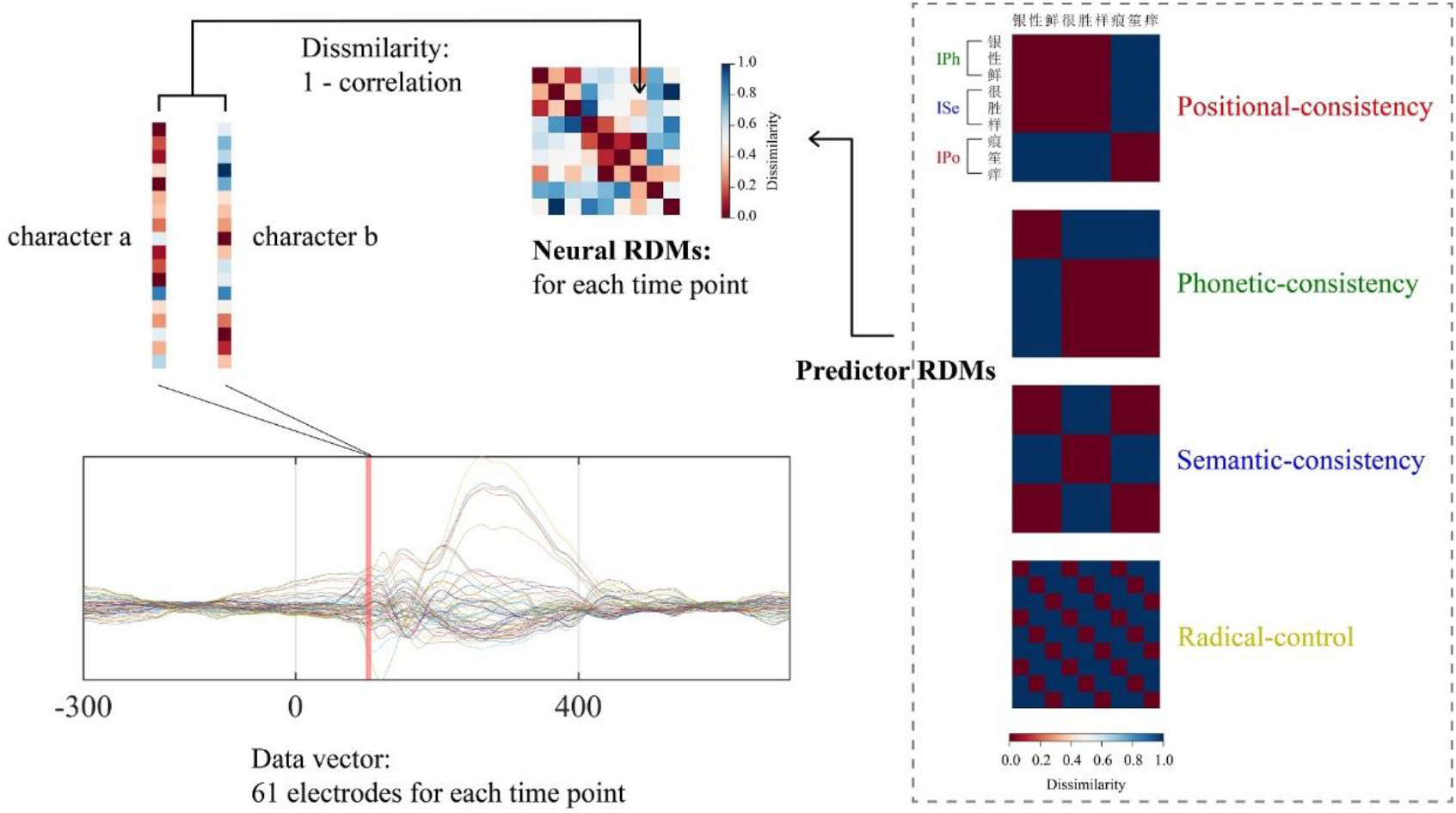
Representational similarity analyses. Neural RDMs are constructed for each data point in grand averaged ERPs by comparing pairwise, phonogram-specific activations. RDMs are symmetric with a diagonal of zeros, and their size corresponds to the number of experimental phonograms, here 9 × 9. Model-based RDMs reflect theoretical predictions. These include three predictor RDMs based on consistency features and one predictor RDM driven by the physical features (phonetic radical types) of the phonograms. After construction, Neural RDMs are compared with predictor RDMs.

#### Modeling neural dissimilarity

In order to characterize the information space that is represented by EEG RDMs, we created three categorical predictor RDMs. Each predictor RDM covered 9 × 9 elements, which were created based on the assumption that there was high similarity within and low similarity across conditions (coded as 0 and 1, respectively). There were three predictor RDMs designed to encode a particular Chinese consistency (Figure 2). For example, we have a positional-consistency RDM (i.e., the dissimilarity between IPh and ISe characters were 0, while the dissimilarity between IPo characters and IPh or ISe characters were 1; the other predictor RDMs have similar rules), a phonetic-consistency RDM, and a semantic-consistency RDM. Furthermore, in order to quantify similarity on the visual features level, we also designed a radical-control RDM based on the phonetic radical category of the material itself (Figure 2). In this RDM, all comparisons that were within the same phonetic radical were marked as similar (0 corresponding to within-category). Additionally, all comparisons between the different phonetic radicals were marked as dissimilar (1 corresponding to between-category). The lower triangles of these predictor RDMs were then correlated (Spearman) with EEG RDMs. These correlations were then computed separately for each time point, which led to a time series of correlations that reflected the correspondence the neural data and the predictor.

#### Statistical Testing

In order to identify significant effects over time, we conducted cluster-based permutation tests (corrected *p* < 0.05) for the complete timeline from stimulus onset to 700 ms. The clusters were formed by five or more neighboring time bins with a cluster-defining threshold of 0.05 (one-sided with the assumption that the correlations between neural RDMs and predictor RDMs were significantly larger than zero) and 1000 iterations (Maris and Oostenveld 2007). For the peaks across time courses, we additionally reported uncorrected t-values and Cohen’s d as a measure of effect size.

### Cross-Temporal RSA

In order to reveal the generalization of representations over time, we constructed RDMs between the neural activities across different time points and calculated the degree of fitting between these cross-temporal RDMs as well as predictor RDMs. The resulting temporal generalization pattern gave information regarding the underlying processing dynamics. For example, a mostly diagonal pattern reflects the sequential processing of specific representations, whereas the generalization from one time point towards another reflects recurrent or sustained activity of a particular process (King and Dehaene, 2014).

The cross-temporal EEG RDMs were obtained by calculating the neural dissimilarity between nine inconsistent characters on one time point, and the nine inconsistent characters on all other time points. The specific calculation settings are consistent within the *neural dissimilarity* section. Therefore, each condition yielded 4900 (70 × 70 time bins) cross-temporal EEG RDMs (9 × 9 elements).

Hence, by calculating the correlations between these cross-temporal EEG RDMs and different predictor RDMs, we were able to investigate the stability of neural representations for various consistencies within their own proprietary time window. In order to identify regions significantly larger than chance level, we conducted a bidimensional cluster-based permutation test on the correlation values that were calculated from each predictor RDM. In this analysis, the adjacent 2 × 2 correlation cubes ere regarded as the smallest cluster, while other parameters were the same as in the previous one-dimensional analysis.

## 3 Results

### Behavioral results

The mean hit rates and mean false alarm rates of the button presses as well as the mean press latencies of correct responses for each experiment are summarized in Supplementary Table 2. The high hit rates (99%) and low false alarm rates (<1%) indicated that the participants were able to accurately focus their attention on the color change detection task.

### vMMNs results

Within each consistency condition, the control and deviant characters evoked the prominent N170-P2-N400 complex at the temporal–occipital recording sites, which was a typical visual ERP (Figure 3). According to a cluster-based permutation test, the vMMN effect was identified for inconsistent positional characters across ROIs in the left and right hemisphere, respectively. The cluster-based permutation test revealed that there was a significantly stronger negativity for the differential wave in left (sum [t] = −383.00; *p* = 0.0128) and right electrodes (sum [t] = −964.46; *p* = 0.0004) for positional-related vMMN. Additionally, a phonetic-related vMMN effect was also found in both electrodes that corresponded to a left cluster (sum [t] = −944.48; *p* = 0.0020) and a right cluster (sum [t] = −869.36; *p* = 0.0016). Moreover, a vMMN effect was also discovered for inconsistent semantic characters in the left cluster (sum [t] = −960.56; *p* = 0.0056) and a right cluster (sum [t] = −303.45; *p* = 0.0290). The time range for significant clusters for various vMMNs across different ROI are shown in Supplementary Table 3.

**Figure 3.**
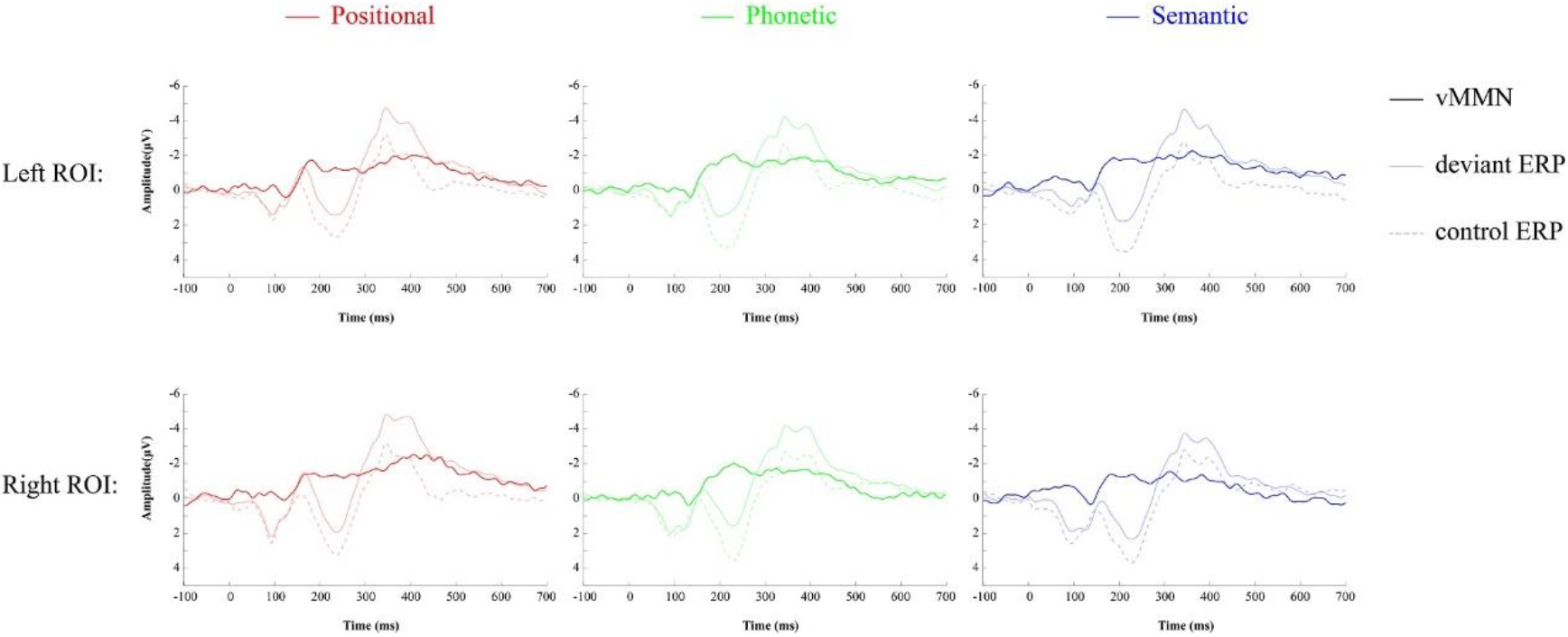
*Light waveform:* Grand-average event-related potentials (ERPs) recorded at left (P7, PO7, O1) and right (P8, PO8, O2) ROI electrodes in response to control (dotted line) and deviant phonograms (solid line) in different dimensions. *Dark waveform:* The vMMN waveforms that obtained by subtracting the standard from the deviant phonograms of each dimension.

### RSA results

In order to reveal positional information in the EEG signals, we generated the positional-consistency predictor RDM, which reflects the nine characters’ dissimilarity in positional consistency dimension. Next, we correlated the neural RDM with this positional-consistency RDM separately at every time bin. This analysis revealed that there was a significant correlation from 160 to 230 ms, peaking at 180 ms (peak t [44] = 2.75 Cohen’s d = 0.83) (Figure 4; red line), which suggests rapidly emerging positional information in the signal. Similarly, we correlated neural RDM with a phonetic-consistency RDM (reflected the nine words’ dissimilarity in phonetic consistency dimension) separately at every time bins and found significant phonetic information from 250 to 520 ms, which peaked at 300 ms (peak t [44] = 2.57 Cohen’s d = 0.77) (Figure 4, green line). Finally, we discovered significant representation (300 to 470 ms) for semantic consistency that peaked at 450 ms (peak t [44] = 2.77 Cohen’s d = 0.84) by correlating the neural RDM with a semantic-consistency RDM. This reflected the characters’ dissimilarity in semantic consistency dimension separately at every time bins (Figure 4; blue line). Additionally, there were, no specific neural representations to radical types found in EEG signals. For example, there was no significant correlation between the neural RDM and radical-control RDM (Figure 4; yellow line). This means that neural activity is not simply driven by sub-lexical features, but rather that it responds to the abstract consistency behind these sub-lexical features.

**Figure 4.**
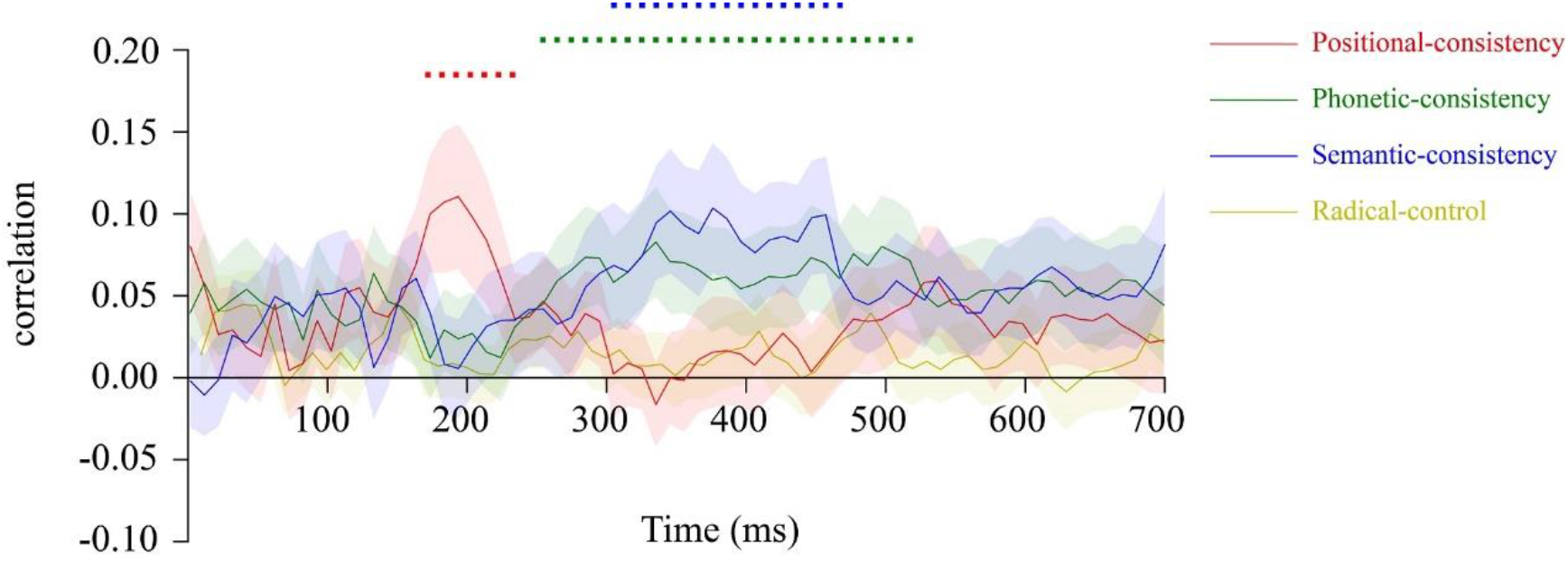
RSA results. We found that the representation time window of positional consistency (red line), phonetic consistency (green line) and semantic consistency (blue line) were respectively emerging between 170 to 230 ms, 260 to 520 ms and 310 to 470 ms. Dots mark significant time windows (cluster-forming threshold *p* < 0.05, corrected for multiple comparisons), and shaded ranges denote standard errors of the mean.

### Cross-Temporal RSA results

We discovered the off-diagonal generalization pattern within the respective time windows of all three orthographic consistency representations. Specifically, the generalization patterns of positional-consistency, phonetic-consistency and semantic-consistency emerged at 160–260 ms, 280–550 ms and 310–470 ms, respectively (Figure 5). It should be noted that a weak but also significant generalization pattern of positional-consistency was also observed in later time periods, around 390–640 ms.

**Figure 5.**
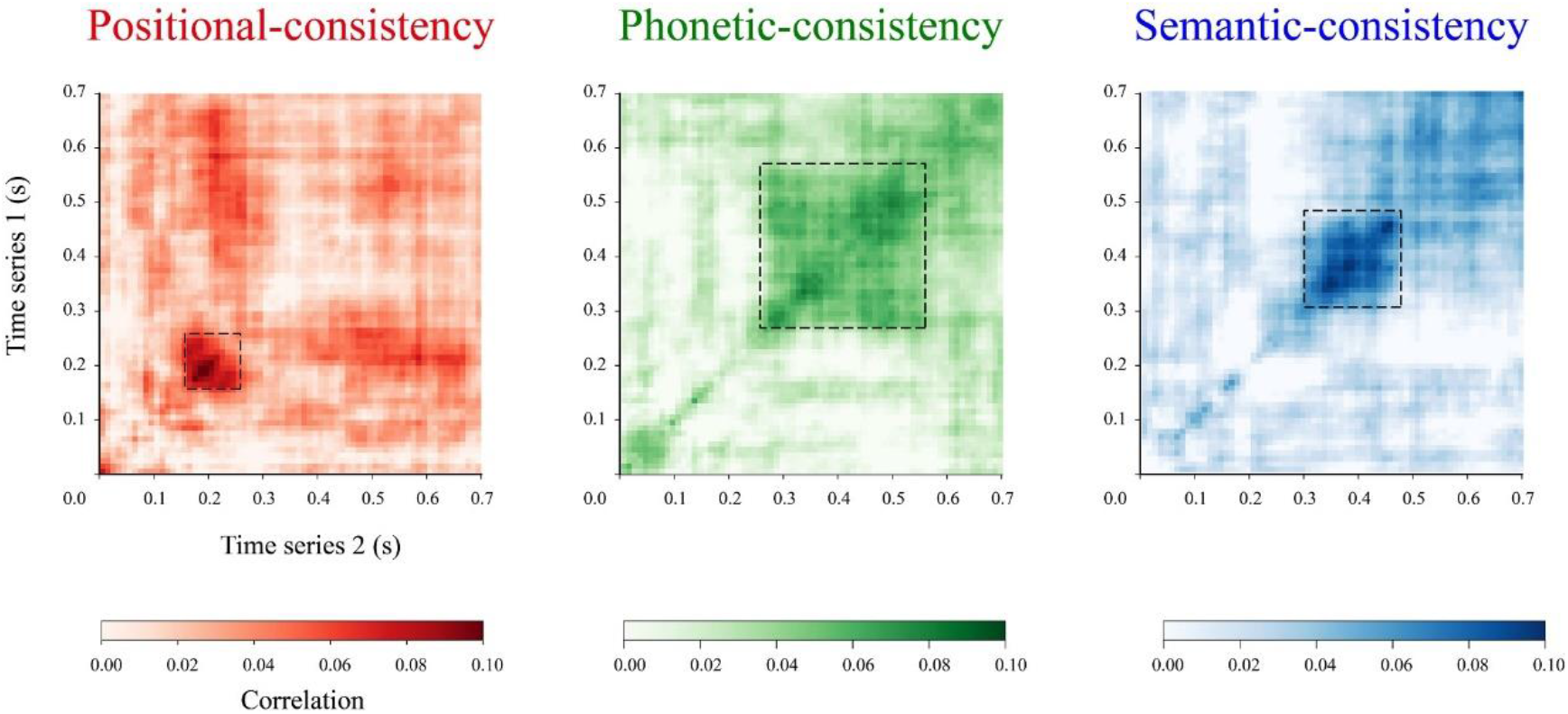
Temporal generalization matrix for each consistency dimension.

## 4 Discussion

The purpose of this study was to investigate the neural mechanisms of automatic consistency processing in visual word recognition. By taking advantage of the rich consistency features that are built on the radical system in Chinese phonograms, we simultaneously manipulate positional, phonetic and semantic consistency across a set of characters. We obtained significant vMMN responses for all three types of inconsistent phonograms, which confirmed the existence of automated electrophysiological activities that were driven by consistency violations. In order to further investigate the temporal dynamic relationship of these automated neural mechanism between the different dimensions, we fitted the neural activities that corresponded to consistency violation with activation patterns that reflected various types of consistency information. These analyses revealed that there was a gradient of distribution between the processing of consistency information on word form and at the lexical level. Interestingly, an off-diagonal generalization pattern is demonstrated in the specific time window of each type of consistency information processing.

One of the important findings in the present study is that the vMMNs that are evoked in the current experiment are best explained by violations of the consistencies category. The vMMN in this study was achieved via subtraction of the ERP to the control characters (in control block) from the ERP to the deviant characters (in oddball block). The control and the deviant characters were comprised of the same phonograms, so that the vMMN cannot be described as physical, orthographical, phonological, or semantic differences between the deviant and the control phonograms. Furthermore, as the probability of presenting the deviant and the control phonograms were exactly equal (i.e., 1/4), then the vMMN cannot be explained as the difference in refractoriness between the ERP to the deviant and the ERP to the control. Moreover, the vMMN cannot be explained as a violation of a phonological category or a semantic category since there is no reducible phonetic or semantic category within the deviant or standard phonograms at our oddball sequence. In summary, through rigorous design, we obtained vMMN responses that reflected the automated handling mechanism of the three types of consistency violations.

More importantly, the automatic mechanism of the vMMN observed in the present study is also sensitive to different categorization process of consistencies. Categorization is a basic process that includes visual perception, and this process goes beyond the physical characteristics of stimulation (Czigler, 2014). For example, there are hundreds of various ‘‘blue’’ colors that have different combinations and values of hue, saturation and brightness. Only when attributed to a dimension of standards and deviants are classified as different types, can the deviants produce surprises (violations) on the predictions that are established by the standard. In the field of phonograms processing, the vMMN effects of word meaning (Hu et al., 2020) and phonological (Wang et al., 2013) categorization have yet to be investigated. The current findings suggest that orthographic consistent or inconsistent features of characters can be placed across different categorical representation systems within the adult brain, not only in orthographic positional categorization, but also in phonological and semantic categorization.

The vMMN response observed in the present study can be interpreted as being the manifestation of a prediction error signal within the predictive coding framework (Friston, 2005; Baldeweg, 2006; Wacongne et al., 2011; Stefanics et al., 2014; Stefanics et al., 2018). In simple terms, predictive coding is the implicit process that creates an internal model regarding sensory input with the aim of minimizing surprise (a quantitative formulation of prediction error, which is the negative log probability of a sensory event) (Friston & Kiebel, 2009). While living in a world of constant change, no two experiences are ever completely the same. Thus, a prediction error will always be present to some degree (Van de Cruys et al., 2014). The predictive coding system, therefore, has to attribute a value or weight to prediction errors in order to determine the extent to which they should induce new learning (a form of meta-learning). A solution to this can be expressed compactly with regards to precision-weighted prediction errors (Friston, 2010), which means that the same prediction violation causes a greater surprise, where predictions are more precise. Back to the current experiment, there has been a lack of effective prediction (low precision) within the control conditions. Furthermore, all characters were decoded using the established neural processing patterns. However, things changed in the oddball condition, where the presence of any unambiguous prediction (high precision) allowed any inconsistent information to be evaluated as worthy of learning, thereby activating any additional neural activity. It makes the same character under the deviant condition evoke a stronger ERP activity compared to the control condition. Thus, we argue that the “genuine” vMMN activities that are evoked by various inconsistent features reflected the high weight processing of the inconsistent phonograms caused by temporally-shaped automatic top-down predictions.

Another important finding of the present study is the temporal dynamics of consistency representation across each dimension under an automatic mechanism. From the beginning of characters present, the neural representation of positional consistency was initially detected, and the peak of these representations occurred around 180 ms. Within the time window of positional consistency representation, neural activity patterns demonstrated a wide generalization, which reflected recurrent or sustained activity of a particular process (not sequential processing) (King and Dehaene, 2014). The time window was consistent with N170/M170 components, which were electric/magnetic activities that were generated by the bilateral occipitotemporal cortex and peak at around 170 ms after the onset of words (Bentin et al., 1999; Tarkiainen et al., 1999; Maurer et al., 2008). These components demonstrated not only selectivity to orthographic stimuli, as it exhibits a more negative amplitude for characters than for other stimuli, but also sensitivity to morphological information of words (Dehaene and Cohen, 2011; Solomyak and Marantz, 2010; Lewis et al., 2011; Fruchter et al., 2013). Based on the available evidence, researchers believe that the reading-related N170 component reflects an early visual expertise in general orthographic processing that distinguishes words (or Chinese phonograms) from other, non-linguistic visual stimuli (Yin et al., 2020). The current findings provide novel insights into the character decoding activities within the time window that correspond to this component, that this stage of cognitive processing is also to distinguish the positional consistency features of phonograms. Future research is required in order to verify whether similar neural representations are able to be detected in word-form consistency in alphabetic characters.

After the time window for positional information, we identified neural representations of phonetic and semantic consistency at 250–520 ms and 300–470 ms, respectively. In addition to the initial time, the peak of phonetic information (300 ms) also appears earlier than semantic information (450 ms). Since consistency representation of the two types of lexical information also demonstrated a wide generalization in their respective time periods, it is reasonable to infer that there is a temporal overlap between the two sustained cognitive activities. The active time window of P200 (around 200–300 ms) and N400 (around 300–500 ms), two typical ERP components of word processing, covered the representation time of phonetic and semantic consistency information. Specifically, the representation activity of phonetic consistency appears to span between two components, while the representation of semantic consistency is concentrated within the time window of N400 component. Based on the two-stage framework, the P200 indexes reflect the variation in the mappings between the orthography and phonology, while the N400 indexes reflect the lexical competition within the selected pronunciations (phonological subgroup). Previous studies that have examined the effect of consistency have captured the reversed responses between the P200 and N400 to consistent and inconsistent speech phonograms. The results have demonstrated that low-consistency characters produce a higher positive amplitude at the P200 time window, and lower negative amplitude at the N400 time window than high-consistency characters (Lee et al., 2007; Hsu et al., 2009). The authors propose that these findings can support the two-stage framework for lexical access, and index the sublexical and lexical processing using P200 and N400, respectively. From this perspective, our results appear to suggest that phonological consistency representations are required within the early stages of sub-lexical processing, and that these representations continue to be effective at the lexical processing stage. In contrast, semantic consistency representations are recruited only during later lexical processing. This also indicates that there is a synergistic effect between phonological and semantic consistency information during the lexical processing stage. This may provide new evidence for interactive theory (Price & Devlin, 2011; Carreiras et al., 2014), and can confirm the contribution of sublexical orthographic information (i.e., O-P and O-S mapping) in order to automatic language processing.

Finally, we should note some limitations of the present study. Several studies have discovered that phonetic consistency of Chinese phonograms is able to also influence N170 amplitude. The results indicated that phonograms with low phonetic consistency evoke a larger N170/M170 than those with high consistency (Lee et al., 2007; Hsu et al., 2009; Hsu et al., 2011). However, we did not find any information that indicated phonetic or semantic consistency in this early time window. This may be due to different phonogram selection methods that were adopted into our study compared to previous studies. Prior studies examined the differences in consistency between the different phonetic radicals, that is, the overall consistency of a certain phonetic radicals were used as a screening criterion for consistent or inconsistent phonograms. While the current study has focused on differences in consistency within the phonetic radicals, a consistent phonogram is a character with a dominant pronunciation of a particular phonetic radical. In fact, our principle fits more closely with the original definition of consistency, which is that competition is among phonological forms of phonograms sharing the same phonetic radical, regardless of the lexical status of the phonetic radical (Yum et al., 2014). Furthermore, there is an important limitation with regards to ERP amplitude differences that are obtained from previous studies of inter-radical consistency. However, the limitation is that, no matter how strictly the additional variables are controlled, there are inevitably some substantial orthographic or lexical information differences between the different radicals, in addition to consistency. This is also the reason why differences in ERP intensity between the different inconsistent conditions were not analyzed in the current study.

## 5 Conclusion

In conclusion, this study discovered consistency violations in visual-orthography, phonetic or semantic dimensions that evoked significant vMMN activities, which indexed an automatic processing mechanism that operates within the predictive coding framework. Moreover, we were able to provide the characterization of the neural dynamics that underlie the automatic processing of different dimensional information. As a whole, these temporal dynamics demonstrated a gradient emergence pattern from orthography to phonology to semantics. However, there remains a certain degree of overlap between phonological and semantic representations, which may indicate the synergy of these two types of information in lexical level processing.

## Acknowledgements

The study was funded by National Natural Science Foundation of China (61807023), Humanities and Social Science Fund of Ministry of Education of the People’s Republic of China (17XJC190010), Shaanxi Province Natural Science Foundation (2018JQ8015), and Fundamental Research Funds for the Central Universities (CN) (GK201702011) to Jingjing Zhao. Queries of the study should correspond to.

## Author contributions

**Jianyi Liu:** Conceptualization; methodology; software; validation; formal analysis; investigation; data curation; writing – original draft; visualization; project administration. **Tengwen Fan:** Investigation; resources. **Yan Chen:** Investigation; data curation. **Jingjing Zhao:** Conceptualization; writing – review & editing; supervision; funding acquisition.

